# Gene set selection for classification with single-cell RNA-seq

**DOI:** 10.64898/2026.11.16.767160

**Authors:** Nikhila P. Swarna, Lior Pachter

**Affiliations:** Division of Biology and Biological Engineering, California Institute of Technology, Pasadena, CA, USA; Department of Computing and Mathematical Sciences, California Institute of Technology, Pasadena, CA, USA

## Abstract

Marker genes are important for analyzing and interpreting single-cell RNA-seq (scRNAseq) experiments. They are used to identify and categorize cell types and can also be useful for the design of downstream experiments. We present CellSieve, which tackles the problem of finding effective gene sets rather than distilling markers one at a time. While several “combinatorial” approaches to gene set selection have been proposed, we find that our approach, which leverages a combination of multinomial logistic regression and linear programming, is much more effective at finding small gene sets that can classify cell types.

## Introduction

The interpretation and utilization of single cell RNA sequencing data is almost always dependent on a cell type annotation. Cell assignments to cell types have many applications, an important one being the identification of “marker genes” for the cell types. Markers serve a dual purpose: they help to develop understanding of the nature and function of cell types, and can also be useful for the design of downstream experiments. The annotation of cell types, usually via clustering, followed by the identification of markers, is part of the standard workflow in the two most widely used single-cell analysis frame-works, Scanpy and Seurat (1, 2). The markers produced are often curated into reference resources that later studies use for cell assignment against (3). Furthermore, marker genes are often used as targets for gene knockouts to study phenotypic effects and biochemical systems in model organisms. For instance, pooled CRISPR screens with single cell readouts have allowed for systematic screens scaling from dozens of perturbed regulators to genome-wide knockdown of over 9,800 genes (4, 5). Marker genes are also used to inform models of gene regulation. Regulon-based methods such as SCENIC use cell type restricted transcription factors as anchors for inferring networks, so the supplied marker gene set directly impacts the recovered network. (6, 7).

Several methods exist to discover marker genes based on distinct and creative ways of formalizing the question in a statistical framework. The most common framing is hypothesis testing, with a comparison of each cell type against all the others, i.e. two-group comparisons (8, 9). Expression-based statistical tests of this kind are the default marker-finding routines in the most widely used single cell toolkits (10). COSG keeps that setting, but replaces the test, constructing an artificial gene expressed only in the target group as the ideal marker and ranking real genes by cosine similarity to it, which is scale independent and scales to datasets of millions of cells (10). The COMET project specifically noted the need for allowing for combinations of genes to mark a cell type, and did so by binarizing each gene with the non-parametric XL-mHG test and returning ranked panels of one or more genes, which it reports as the first prediction of multi-gene marker panels ranked by relevance (11). NS-Forest instead frames the task as random forest feature selection followed by binary expression scoring, and it discovers minimal marker gene combinations (12). scGeneFit frames it as label-aware compressive classification and optimizes joint label recovery, which yields markedly less redundant panels than one-versus-rest selection (13). geneBasis drops labels entirely and adds genes iteratively, so that each new gene captures the maximum distance between the true manifold and the manifold built from the currently selected panel (14). A recent benchmark assembled 59 computational methods for marker selection and evaluated them on 14 real and over 170 simulated datasets (15). Several tools exist for downstream analyses based on selected markers as well. Standardized pipelines now wrap marker identification together with its evaluation (16). Probe set selection for targeted spatial assays has become its own problem, and the methods built for it deliberately move past marker-driven selection in order to capture continuous spatial signal and new states (17, 18).

The term marker gene is often used as a blanket term for highly variable genes, highly specific genes, or differentially expressed genes, but it is important to recognize how these three definitions interact to form an effective marker panel. Highly variable gene selection targets cell to cell variance within a population and is label free by construction. Specificity scores such as *τ* and Ψ measure how concentrated expression is across partitions of a given data set, and the scores are directly influenced by how cells are annotated (19, 20). Differential expression measures how a gene’s expression varies between pairwise conditions using contrasts (21). Depending on the biological context and the target information to be extracted from a dataset, a combination of these definitions can form the most effective marker panel.

In this paper we pose a specific question: What is the smallest set of marker genes needed to accurately recover cell type annotations in a dataset? This question is of special importance for spatial transcriptomics, where panel selection requires marker genes that are dense with classification information, and that can translate across datasets and modalities (18, 22, 23). Targeted spatial panels typically cover less than one percent of the genome (18). A massive whole-brain MERFISH atlas imaged just over 1,100 genes to resolve more than 5,000 clusters across roughly 10 million cells (24), using a panel designed by selecting the genes with the greatest combinatorial power to discriminate among all clusters (25). Each added HCR target costs a separate spectrally distinct amplifier channel, so panel size is a hard experimental budget rather than a soft preference (26).

Our approach is a two-stage process, where we first model cell type classification using multinomial logistic regression and then use this model to inform a mixed integer linear programming optimization problem to find an small set of genes needed to compress the given classification. We demonstrate that our approach results in increased classification accuracy across panel sizes using a multi-tissue dataset of 5.2 million nuclei spanning eight tissues, eight founder strains and 106 cell types and states (27). We discover that minimal cell type markers are highly combinatorial and carry biological signal that is effective at translating from sequencing experiments to spatial experiments (24, 27). We show-case that modeling cell type in a dataset using minimal combinatorial panels can refine and correct mistakes in existing cell type annotations in the Zhengmix4eq data (28).

## Results

### A multinomial logistic regression prior to an MILP produces competitive and compact marker gene panels

CellSieve uses a two stage approach to find minimal marker panels. It first sieves through thousands of genes to distill the data down to compact sets that preserve the most classification accuracy within a multinomial logistic regression model with an elastic net prior (29) to model cell type classification in a given dataset. The fit relieves the optimization bottle-neck by prioritizing genes that carry the most classification information along with the cells that are most discriminative of cell type (*Methods*).

We then frame the search for the minimal gene set, given a fixed annotation, as a mixed integer linear program (MILP). The objective minimizes weighted classification slack while maximizing selection of the genes favored by the elastic net prior. CellSieve also offers the option to find interchangeable solutions, meaning alternative panels whose genes serve similar combinatorial roles in the resulting classification problem (Fig. 1a).

To test how CellSieve performs compared to existing marker panel extraction methods, we benchmarked CellSieve in C57BL/6J (B6J) Cortex/Hippocampus tissue from the Rebboah et al. 2026 data (27). The benchmark dataset contains 86,427 cells, 23 annotated cell types, and 5,083 candidate genes that were preselected using ember to be highly informative by the cell type partition (20). We compared against scGeneFit (13), panels adapted from native NS-Forest rankings (12, 30), and L1 (lasso) and L2 (ridge) logistic-regression coefficient rankings (31). We evaluated gene panel performance using a common class balanced logistic regression classifier and found that CellSieve produced panels with higher macro-F1 recovery than its competitors across all panel sizes (Fig. 1b).

**Fig 1.**
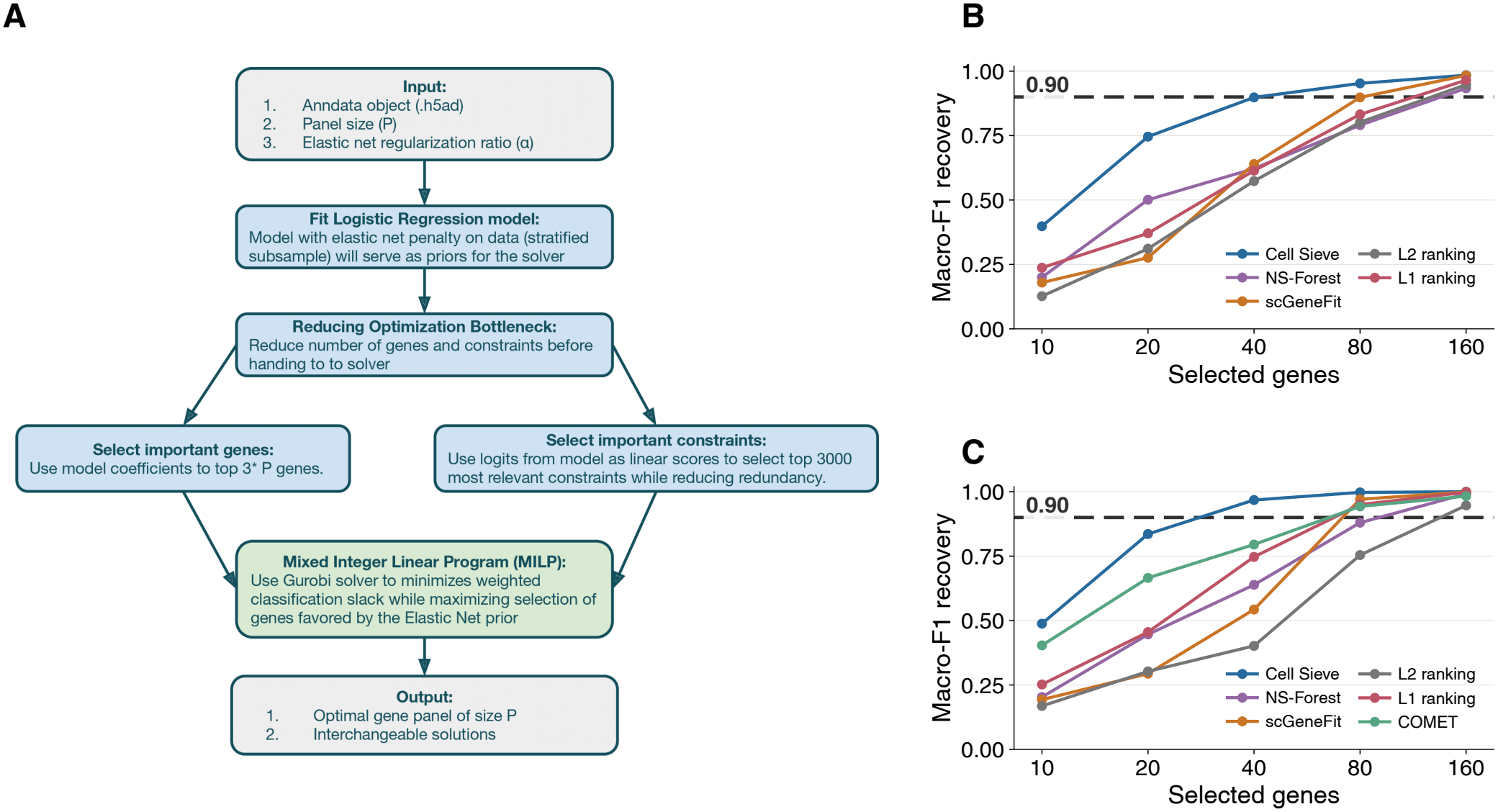
CellSieve selects compact gene panels that recover cell-type annotations. **a**. Overview of the CellSieve workflow. Elastic-net logistic regression provides gene and constraint priorities for a mixed-integer linear program. CellSieve uses a Gurobi solver to extract an small gene panel with alternative gene swaps. **b**. Benchmark on the full B6J cortex dataset, comprising 86,427 cells, 5,083 genes, and 23 annotated cell types. CellSieve is compared with NS-Forest, scGeneFit, and gene rankings derived from L1- and L2-regularized logistic regression. **c**. Benchmark on a common, cell-type-stratified subset of 2,500 cells from the same dataset, retaining all 5,083 genes and 23 cell types. The same methods are evaluated, with COMET included in this smaller benchmark. In both benchmarks, panels contain 10, 20, 40, 80, or 160 selected genes. Curves show macro-F1 for recovery of the reference cell-type annotations after classifier refitting and evaluation on the same benchmark cells. Dashed lines indicate a macro-F1 of 0.90.

It is particularly interesting that converting multinomial logistic regression classification into an optimization problem allows for gene panels that significantly outperform pure lasso and ridge regression. While multinomial logistic regression is a powerful tool to model cell type classification, it struggles to prioritize genes to produce compact gene panels, making CellSieve’s optimization framework a critical addition.

To include COMET (11) under its cell-count constraint, we repeated the same comparison on a common stratified subset of 2,500 cells retaining all 23 types and the same genes. CellSieve achieved the highest macro-F1 at every panel size even in this subset (Fig. 1c).

The complete benchmark readouts are provided in the supplement, including accuracy, three-fold classifier cross-validation of macro-F1 and accuracy, wall clock run time and memory usage (Supplementary Fig. S1). While CellSieve requires higher memory and runtime than its competitors, it provides superior classification accuracy and macro-f1.

We additionally extend this benchmark against the tools scGeneFit and NS-Forest to all 8 tissues in the Rebboah et al. data within B6J and to all strains within the Cortex Hip-pocampus (Supplementary Fig. S2-4). Across tissues, strains and panel sizes CellSieve has higher macro-F1 and accuracy on average. These results showcase the effectiveness of CellSieve’s extension of multinomial logistic regression using an optimization framework across contexts and present it as a state of the art and competitive marker panel extraction tool.

### CellSieve produces combinatorial gene panels that are interpretable and transfer across modalities

Compressed representations of structure in single cell data are usually latent variables, learned by variational autoencoders or neural networks (32, 33). These methods are both more computationally expensive and do not allow for clear interpretation of resultant gene panels. CellSieve’s gene panels can be interpreted directly by fitting logistic regression models based on the resultant panels and studying their coefficients.

We can inspect the 40 gene panel that passed the 0.9 macro-F1 threshold from Figure 1b and study its model coefficients to learn how each of the 40 genes is contributing to the classification task (Fig. 2a).

**Fig 2.**
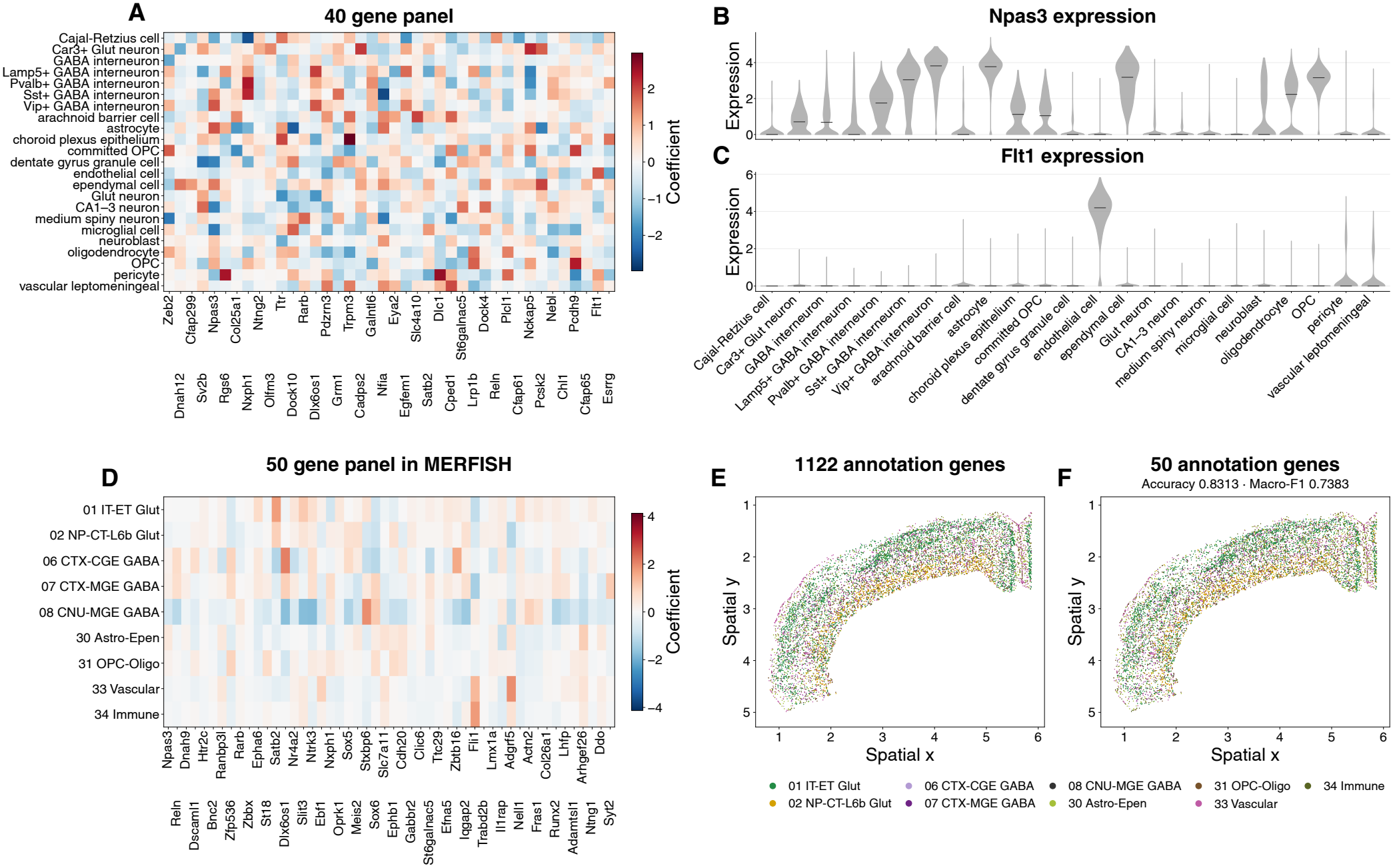
Interpreting Cortex marker panels generated by CellSieve. **a**. Multinomial logistic regression model coefficients from the 40 gene panel generated in Fig 1b **b–c**. Gene expression of *Npas3* and *Flt1* across all 86,427 cells, grouped by cell type; black marks indicate medians. **d**. Multinomial logistic regression coefficients for a 50-gene panel selected from the overlap genes between Rebboah et al. 2026 8cube cortex hippocampus in B6J (27) and Zhang et al. 2023 MERFISH spatial cortex data (24); fit to the Zhang et al. 2023 cells. **e**. Original Zhang et al. 2023 cell type annotations **f**. Cell type annotation prediction for 8,891 cells using 50 gene panel in d. Accuracy is 0.8313 and macro-F1 is 0.7383.

Examining two genes closely, *Npas3* and *Flt1* we can see that marker panels can consist of genes that are both highly variable and highly specific. *Flt1* shows exclusive expression in endothelial cells within the B6J cortex and is a well recognized marker for this cell type within the cortex (Fig. 2c) (34). *Npas3* shows expression across several cell types in the cortex (Fig. 2b) and has been documented to play a critical role in murine development across developmental stages and tissues (35). Its expression spans 11.5 to 17.5 days post coitum and extends to non-neural tissues such as the developing dermis and mesenchyme, and mice lacking it lose 84% of neural precursor proliferation in the dentate gyrus (36).

Furthermore we did a GO analysis on the 40 gene panel (Table 1) and found that several selected genes have developmental annotations. This finding is in line with our understanding of how cell types are formed through the process of differentiation (37).

To further test how biologically relevant CellSieve selected genes may be, we performed an analysis to see how well cell type classification accuracy and macro-F1 transfer to unseen datasets. We first took the overlap of genes between Rebboah et al. 2026 snRNA-seq B6J cortex and the Zhang et al. 2023 MERFISH cortex atlas and used the overlap genes to generate a 50 gene marker panel in the snRNA-seq data (Fig. 2d). We then evaluated the sequenceing selected panel using cross-validation within the MER-FISH data: we achieved an 0.8313 accuracy and a 0.7383 macro-F1 (Fig. 2e,f). These findings show that CellSieve panels are uniquely interpretable and carry latent signal encoding cell type processes that transfer across datasets and modalities.

### Combinatorially compressed marker panels can refine existing cell type annotations

Upon learning that CellSieve is able to identify genes that carry meaningful biological signal encoding cell type, a natural follow up question is whether such a framework can assist with cell type annotation. So we first engineered an experiment within the same Rebboah et al. 2026 B6J cortex data in which we corrupted 10% of the cells with random noise and asked if an 80 gene CellSieve marker panel generated from the corrupted data could identify cell type annotation errors.

The 80 gene CellSieve model was able to identify 5,972 of the 8,642 intentionally mislabeled cells and assign them back to their correct cell type. Only 321 were changed to an incorrect label and 2,349 cells were missed. In addition to recovering injected noise, the model also identified 1,251 non-corrupted cells as misannotated and changed them away from the reference (Fig. 3a).

**Fig 3.**
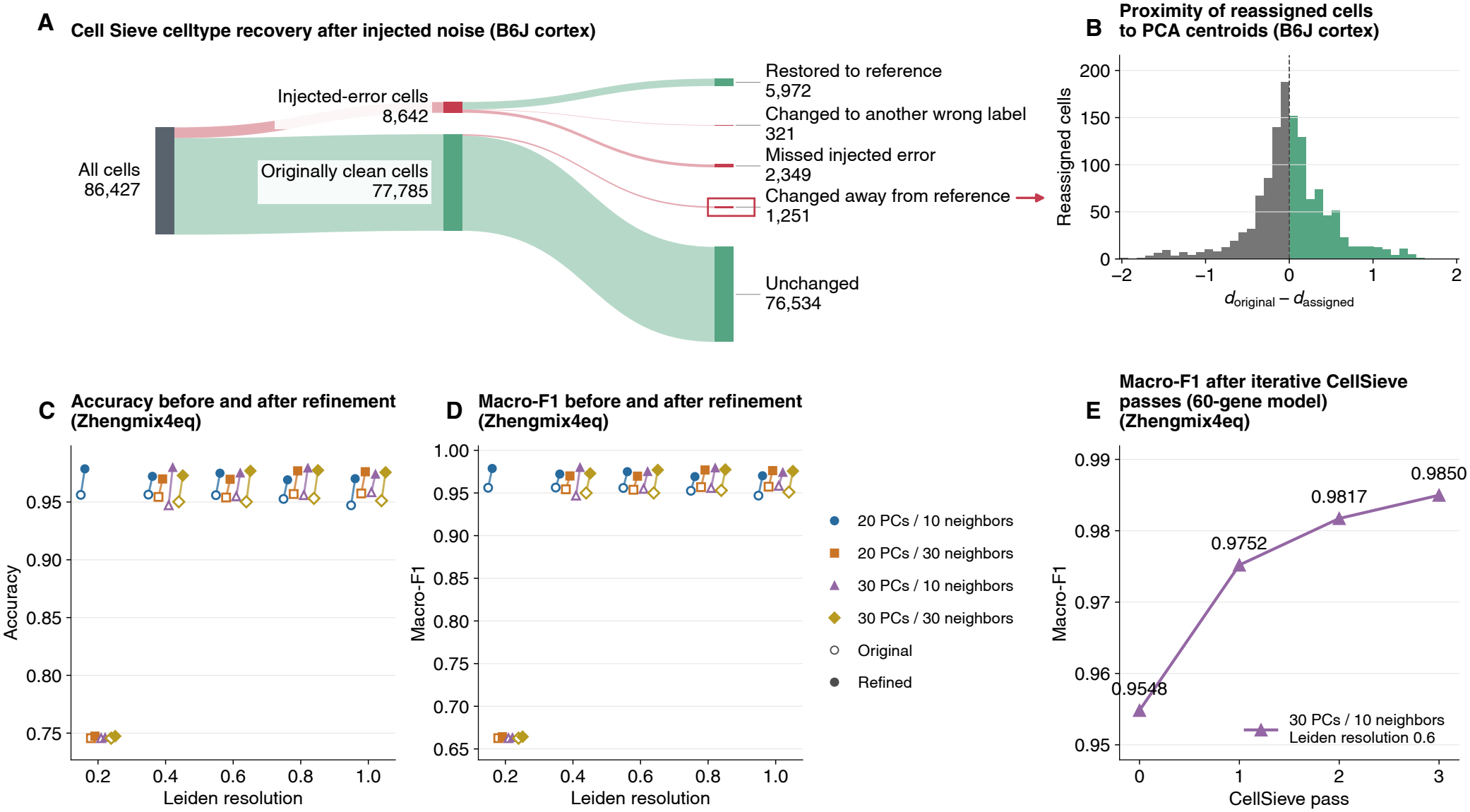
Demonstrating cell type annotation refinement using CellSieve. **a**. Rebboah et al. 2026 cortex data injected with 10% random noise and following recovery using one 80-gene CellSieve pass restored 5,972 of 8,642 injected errors; 321 changed incorrectly and 2,349 were unchanged. Of 77,785 unperturbed cells, 1,251 changed away from the reference. Groupings of cells with labels aligned with the original reference are colored green throughout the alluvial plot and those mislabeled are in red. **b**. Distance to the original median center minus distance to the assigned center across 50 PCs for those 1,251 cells. Positive values colored green favor the assigned type (620 cells); negative values colored gray favor the original type (631 cells). **c–d**. Changes in accuracy and macro-F1 across 20 separate unsupervised clustering instances of the Zhengmix4eq (28) data with different initialization settings. Each clustering is refined once using CellSieve with 60 gene panels. **e** Accuracy and macro-F1 over three CellSieve passes for 30 PCs, 10 neighbors, and resolution 0.6.

We further inspected these 1,251 cells to see if they could in fact be misannotation in the data that we are treating as ground truth, but is not necessarily so. Cell type annotation is an inherently difficult process that requires iteration and biological knowledge when it isn’t paired with some cell sorting mechanism (38). We used the first 50 PCs to measure the distance of these cells from their original and misassigned cell type label centroids in PCA space and discovered that 620 of the 1,251 cells were closer to the CellSieve annotation than their originally assigned one (Fig. 3b). The remaining 631 sat closer to their original label, so the split is close to even and we read a disagreement as a prompt for review rather than as proof that the reference is wrong. While this corruption experiment serves as a proof of concept that blatant mistakes can be caught in an engineered experiment, how well does CellSieve perform in an unsupervised setting where cell type labels are unknown?

We next took the very popular Zhengmix4eq dataset that is often used for benchmarking clustering and cell type assignment pipelines due to its distinct experimental design. 4 FACS sorted cell types that are sequenced along with their ground truth cell type labels. Since the question is how well does CellSieve improve cell type annotations in a realistic setting, we first clustered this data using Leiden clustering with 20 different initialization settings: 20 or 30 PCs, 10 or 30 neighbours and leiden resolution ranging from 0.2 -1.0. In each of these settings we applied a single CellSieve refinement by fitting a 60 gene model to the resulting clusters and evaluating how well these clusters mapped back to existing FACS annotations. Across all 20 conditions CellSieve refinement improved both macro-F1 and accuracy (Fig. 3c,d).

In the condition with highest macro-F1 before refinement, 30 PCs, 10 neighbours and 0.6 Leiden resolution, we did 3 iterative cell type annotation refinements using CellSieve and witnessed a jump in macro-F1 from 0.9548 to 0.9850 (Fig. 3e). Each round provided incremental improvements.

These results showcase CellSieve’s ability to learn cell type classification and improve annotations, serving as a tool that can be easily and reliably integrated into existing cell type annotation pipelines.

## Discussion

A major takeaway from our findings is that a marker gene is highly context dependent. It is a property of a gene, a cell type, and the set of alternatives being considered. The same gene can be a textbook marker in one tissue and uninformative in another. Consider our example where *Flt1* separates endothelial cells from other cells on its own, while *Npas3* is graded across cell types and only becomes informative in combination with other genes. The context dependence of marker genes has been widely noted and studied (20, 39). This context dependence also explains why 59 published marker selection methods can disagree while each performs well under its own definition (15). This motivates a careful interpretation of curated marker databases as a record of what has been observed rather than as a fixed ground truth (3). CellSieve is a tool built on this vital framing that any marker gene is highly context dependent, and so we define a marker panel entirely by what it can classify, given one annotation.

While CellSieve employs a rigorous multinomial logistic regression classifier paired with a mixed integer linear program, it should be noted that this tool employs several heuristics to provide timely and compute efficient solutions. The elastic net prior prunes the gene universe to a candidate set before the optimizer ever runs, so a gene discarded at that stage cannot be recovered later. We solve the resulting program to a relative gap target rather than to optimality, under a wall clock limit. In practice, the larger full-cortex panels exhaust that limit, and the 40, 80 and 160 gene panels finished with reported gaps of 63.03%, 97.56% and 104.31%. Those panels are feasible incumbents, not certified global optima. The panels still outperform the comparators at small budgets, which tells us the formulation is useful, not that we have found the smallest possible panel.

The optimization bottleneck here is not just an implementation detail and is worth discussing further. Choosing a smallest set of genes that distinguishes every pair of cell types can be framed as a set cover problem: take the pairs of cell types as the ground set, and let each gene cover the pairs it separates. Set cover sits among the original NP-complete problems (40), and it cannot be approximated to better than a factor of ln *n* unless P equals NP (41), a bound the simple greedy algorithm already attains (42). A set cover framing of marker selection would be attractive because it states the combinatorial requirement directly, with one constraint per cell type pair. This is however, a very expensive framing, since the number of pair constraints grows quadratically in the number of types, and the number of binary variables grows with the gene universe. A hard cover formulation is also fragile on single cell data. Whether a gene separates a given pair of cell types has to be committed as a binary fact, and that commitment relies on counts shaped by dropouts and uneven sequencing depth (43–45). A gene that covers a pair at one depth may fail to cover it at another, so the constraint set is not stable across datasets. This is why CellSieve pays for classification violations with weighted slack rather than demanding exact coverage. Our constraint sampling and candidate pruning exist to keep that program tractable, and both are deliberate approximations. We do not expect exact minimal panels to be reachable at atlas scale. While an exact formulation would be satisfying, it risks solving the wrong problem. Single cell counts are sparse and unevenly sampled, and there is no external ground truth against which a panel can be validated. When the objective is defined against labels we cannot independently verify, heuristics are an appropriate response to a high dimensional problem whose optimum is only as meaningful as the annotation it is measured against.

A related limitation of our framework is that since CellSieve compresses against an annotation, it inherits whatever that annotation gets wrong, and fine-grained cell types are exactly where annotations are least stable. Cluster assignments are are often hard to replicate and change with the number of principal components, the neighborhood size and the resolution parameters used (28, 46). While our frame-work allows for annotation refinement to a certain degree, large mistakes in annotation will bleed into resultant classification models. A discriminative model needs a label set to be handed to it (47), which makes it poorly suited for deciding how many types there are in the first place. Generative models of expression take the opposite stance and learn a latent representation without committing to a fixed taxonomy (32). Pairing a generative model of the data with a discriminative compression step is the natural next design, and it would let panel selection and type definition inform each other rather than running in series.

**Table 1.** Gene ontology analysis for selected 40 gene panel in Rebboah et al. 2026 mouse cortex.

| GO ID | Biological process | Raw *P* | Selected genes |
|---|---|---|---|
| GO:0007399 | nervous system **development** | 3.05 × 10<sup>−5</sup> | *Chl1*, *Col25a1*, *Dlc1*, *Dock10*, *Nfia*, *Ntng2*, *Olfm3*, *Pcsk2*, *Rarb*, *Reln*, *Satb2*, *Slc4a10*, *Zeb2* |
| GO:0048731 | system **development** | 4.965 × 10<sup>−4</sup> | *Chl1*, *Col25a1*, *Dlc1*, *Dock10*, *Flt1*, *Nebl*, *Nfia*, *Ntng2*, *Olfm3*, *Pcsk2*, *Rarb*, *Reln*, *Satb2*, *Slc4a10*, *Zeb2* |
| GO:0048856 | anatomical structure **development** | 1.835 × 10<sup>−4</sup> | *Cfap61*, *Cfap65*, *Chl1*, *Col25a1*, *Dlc1*, *Dnah12*, *Dock10*, *Eya2*, *Flt1*, *Lrp1b*, *Nebl*, *Nfia*, *Ntng2*, *Olfm3*, *Pcsk2*, *Rarb*, *Reln*, *Satb2*, *Slc4a10*, *Zeb2* |
| GO:0007275 | multicellular organism **development** | 5.991 × 10<sup>−4</sup> | *Chl1*, *Col25a1*, *Dlc1*, *Dock10*, *Flt1*, *Lrp1b*, *Nebl*, *Nfia*, *Ntng2*, *Olfm3*, *Pcsk2*, *Rarb*, *Reln*, *Satb2*, *Slc4a10*, *Zeb2* |
| GO:0032502 | **developmental** process | 4.995 × 10<sup>−4</sup> | *Cfap61*, *Cfap65*, *Chl1*, *Col25a1*, *Dlc1*, *Dnah12*, *Dock10*, *Eya2*, *Flt1*, *Lrp1b*, *Nebl*, *Nfia*, *Ntng2*, *Olfm3*, *Pcsk2*, *Rarb*, *Reln*, *Satb2*, *Slc4a10*, *Zeb2* |

The question we would most like to answer is one this work does not settle: given a dataset and an annotation, what is the absolute smallest number of genes that could recover the labels? A per-dataset lower bound would turn panel design from a search into a budget, and it would tell an experimentalist when adding probes has stopped buying information. The difficulty is that the bound depends on measurement as much as on biology. Several statistical frameworks already incorporate technical noise into models of gene expression, including multinomial sampling models (43), probabilistic models of capture efficiency and sequencing depth (32), and more recent approaches that distinguish biological variation from technical variation in spatial transcriptomics (48). Building on these approaches, a meaningful direction for future work would be to develop a multivariate classification framework that treats sequencing depth, capture efficiency, and the resulting dropout probabilities as explicit, tunable variables alongside the biological classification objective. Such a model could quantify how the minimum number of genes required to recover a given annotation changes with measurement quality, providing a direct relationship between classification performance, experimental design, and the information density of a dataset. This would allow gene-panel selection to inform not only which genes should be measured, but also how deeply they must be measured to preserve the biological information needed for classification. We think this is a meaningful direction for future work as it would allow for a quantitative link between panel performance and data quality, making biologically informed data generation and harmonization more tractable.

## Methods

### CellSieve algorithm

#### Logistic fitting and global approximation

CellSieve first estimates gene informativeness using an elastic-net logistic classifier. To reduce fitting cost, this initial model uses stochastic gradient descent with one-versus-rest log loss on at most 15,000 stratified, maximum-absolute-value-scaled cells. This approximates a full-data joint multinomial fit. The elastic-net L1 ratio *α* controls sparsity, while balanced class weights account for unequal cell-type abundance. Each gene receives a prior score *q*_*g*_ equal to its largest absolute coefficient across classes.

#### Reducing the optimization bottleneck

For a panel of *k* genes, CellSieve restricts optimization to the top min(4*k*, |*G*|) prior-ranked genes, where *G* is the retained gene pool. Balanced L2 multinomial logistic regression is refitted on standardized candidate expression. Its coefficients remain fixed during gene selection, avoiding joint optimization of classifier parameters and panel membership.

The algorithm further reduces the problem to 3,000 cell/label comparisons, selected using classification margins and penalties for repeated cells and label pairs. For comparison *s* = (*i, t, w*) between cell *i*’s assigned label *t* and comparison label *w*, define

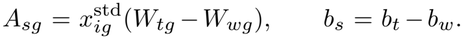

These quantities describe each gene’s contribution and the intercept contribution to the reference classifier’s label-score difference.

#### Objective and hyperparameters

CellSieve selects exactly *k* genes by solving

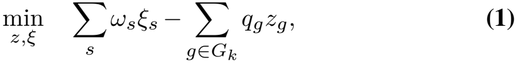

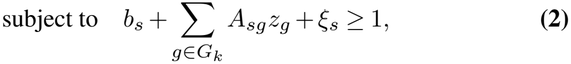

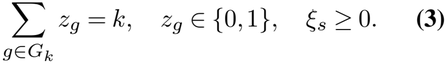

Here *z*_*g*_ indicates gene selection and *ξ*_*s*_ measures a margin shortfall. Weights *ω*_*s*_ balance sampled label pairs and increase penalties for less abundant assigned classes. The objective rewards informative genes while penalizing lost separation.

The global fit uses regularization strength 0.001, at most 1,000 iterations, tolerance 10^*−*3^, and seed 42 (31). Gurobi (49) uses four workers and relative gap target 0.10, with time limits of 600 seconds for full-cortex and overlap-constrained panels and 300 seconds for subset and supplemental panels. It should be noted that time-limited feasible solutions are not certified global optima.

#### Benchmarking methodology

We benchmarked genepanel selection on a B6J mouse cortex dataset (27) containing 86,427 cells and 23 annotated cell types. Before benchmarking, genes were filtered using ember (20) with cell type as the partition, retaining genes with significant overall specificity (*p*_Ψ_ *<* 0.05) and Ψ *>* 0.75, and excluding genes with `Gm`, `mt-`, `Rps`, `Rpl`, or `ENSMUSG` prefixes or a `Rik` suffix. This yielded a common search space of 5,083 genes to accommodate slower L1 ranking baseline. We first evaluated all cells and then a common stratified subset of 2,500 cells, retaining all 23 cell types. The panel sizes examined were 10, 20, 40, 80, and 160 genes.

#### CellSieve

CellSieve was run with an elastic-net L1 ratio of 0.7. The global prior was fitted on at most 15,000 stratified cells using maximum-absolute-value scaling. Candidate genes were prioritized by their elastic-net coefficients, followed by localized logistic-regression refitting and mixedinteger optimization. Gurobi (49) used a relative optimality-gap target of 0.10 and time limits of 600 seconds per panel for the full cohort and 300 seconds per panel for the subset.

#### NS-Forest

NS-Forest (12, 30) was run through its native API, including median-expression and binary-score preprocessing, with 160 candidate genes and 160 binary genes per cell type. To obtain panels matching the requested gene budgets, class-specific binary-gene rankings were combined by within-class rank, with ties resolved by descending binary score and gene name. Duplicate genes were removed, and the first *k* genes were selected. This comparison used panels adapted from NS-Forest’s native ranked outputs.

#### scGeneFit

scGeneFit (13) was run using its native `get_markers` function with the `centers` method on standardized expression, requesting the specified number of genes.

#### COMET

COMET (11) was evaluated on the common 2,500-cell subset because its native cell-count limit precluded the full cohort. Native singleton rankings were combined across classes by rank, with ties resolved by ascending minimum-hypergeometric statistic and gene name. Negated markers were mapped to their measured genes, duplicate genes were removed, and the first *k* genes were selected.

#### L1 and L2 ranking baselines

Genes were ranked by the maximum absolute logistic-regression coefficient across cell types, and the top *k* genes were selected. L1 ranking used class-balanced logistic regression with the `saga` solver (31, 50). L2 ranking used class-balanced logistic regression with the default L2 penalty.

#### Refit and cross-validation evaluation

Every selected panel was evaluated using the same class-balanced logisticregression classifier (31), with a maximum of 2,000 iterations. For refit evaluation, selected-gene expression was standardized across the benchmark cohort, and the classifier was fitted and evaluated on those same cells. Accuracy was the fraction of correctly classified cells. Macro-F1 was the unweighted mean of cell-type-specific F1 scores:

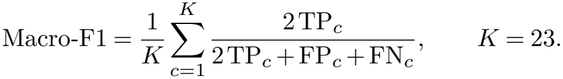

Undefined class-specific F1 scores were assigned zero.

We additionally performed shuffled, stratified three-fold cross-validation with random seed equal to 42. Scaling and classifier fitting used only the training cells within each fold. Accuracy and macro-F1 were calculated on held-out cells and averaged across folds. Gene panels were selected once using the complete benchmark cohort and held fixed during cross-validation. These CV scores therefore measure conditional classifier performance for an already selected panel. Figure 1 reports refit macro-F1; the supplementary figure reports conditional CV macro-F1, refit and CV accuracy, and resource measurements.

#### Supplemental Benchmarks

Supplemental benchmarks compared CellSieve, NS-Forest, and scGeneFit across nine B6J tissue datasets (27) (adrenal, cortex/hippocampus, diencephalon/pituitary, gastrocnemius, female gonads, male gonads, heart, kidney, and liver) and, separately, across eight founder strains (129S1J, AJ, B6J, CASTJ, NODJ, NZOJ, PWKJ, and WSBJ) (51) in cortex/hippocampus. Each tissue– strain cohort was benchmarked independently using its cell-type annotations. Cell types represented by at least 10 cells were retained, and genes with `Gm`, `mt-`, `Rps`, `Rpl`, or `ENSMUSG` prefixes or a `Rik` suffix were excluded. CellSieve used tissue-specific elastic-net L1 ratios selected as described below and a 300-second solver time limit per panel. Evaluation followed the same procedure as the main figure, including refit and conditional three-fold CV macro-F1 and accuracy, wall-clock runtime, and peak process-tree memory.

For each B6J tissue, the elastic-net L1 ratio was selected from *α* ∈ {0.1, 0.3, 0.5, 0.7, 0.9} using 20 stratified subsamples of 5,000 cells, retaining at least 10 cells per cell type. Within each repeat, all *α* values shared the same approximately 75%/25% stratified training–test split. CellSieve selected a 40-gene panel using training cells only, and a class-balanced logistic regression classifier was evaluated by held-out macro-F1. Differences among *α* values were assessed using a within-repeat permutation test of the Friedman rank statistic and exact two-sided paired sign-flip tests, with Benjamini–Hochberg correction across the ten pairwise comparisons within each tissue. The *α* with the highest mean held-out macro-F1 was selected and used across panel sizes: *α* = 0.7 for adrenal, diencephalon/pituitary, and kidney; *α* = 0.5 for heart; and *α* = 0.9 for the remaining tissues. All eight cortex/hippocampus strain benchmarks used *α* = 0.9. No tissue had a unique winner supported against every alternative after multiplicity correction; these selections therefore represent empirical performance maxima within the tested grid. (Supplementary Fig. S2-4)

#### Gene Ontology analysis

We performed Gene Ontology (GO) over-representation analysis on the 40-gene cortex marker panel using mouse annotations from Mouse Genome Informatics. The background comprised 25,053 mouse genes with positive, non-root GO annotations. Of the 40 selected genes, 38 had usable annotations; *Dlx6os1* and *Otx2os1* were excluded. Annotations were propagated through `is_a` and `part_of` relationships, retaining electronic annotations and excluding negative (`NOT`) and no-data (`ND`) annotations. Enrichment was assessed using one-sided hypergeometric tests, with Benjamini–Hochberg correction across 21,889 biological process, molecular function, and cellular component terms. The table reports raw, unadjusted *P* values for five selected developmental biological processes; no tested term met an FDR threshold of 0.05.

#### Cortex expression and MERFISH

Cortex coefficients and violin plots use the native 40-gene Figure 1 panel. Violin plots use all cells and unmodified AnnData (52) `X`, without an additional transformation. The 50-gene panel was selected independently from the gene-name intersection, retaining cortex gene order and values.

MERFISH values (24, 25) were marked `log2p` and were not logged again. Cell types with fewer than 10 cells were excluded. Balanced L2 logistic regression used *C* = 1, `lbfgs`, at most 2,000 iterations, tolerance 10^*−*4^, and seed 42. Five shuffled stratified folds used seed 42, with standardization fitted within each training fold.

#### Annotation refinement

Native five-fold probabilities changed labels only when the highest-probability class differed from the assigned class and exceeded its probability by more than 0.30. Native selection and scaling precede classifier CV, so predictions are not fully nested validation.

Cortex noise injection uniformly sampled 8,642 cells without replacement with seed 42, assigning each a random label other than its reference. One 80-gene, L1-ratio-0.7 pass was evaluated. For proximity, raw counts were normalized to 10^4^, `log1p`-transformed, and scaled with clipping at 10 using the original highly variable gene mask (1). Fifty PCs used ARPACK (53) with seed 0. Centers were median vectors of unperturbed, unflagged cells, requiring at least 20 per type. Distances use all 50 PCs.

For Zhengmix4eq (28), genes expressed in at least three cells were retained. Counts were normalized to 10^4^ and `log1p`-transformed. Initializations used 2,000 Seurat highly variable genes (1, 54), scaling clipped at 10, 50-PC ARPACK PCA (53), exact sklearn neighbors (31), and native Leiden with seed 42 (55). The grid varied PCs (20/30), neighbors (10/30), and resolution (0.2/0.4/0.6/0.8/1.0). Frozen matrices and partitions were reused. CellSieve used all retained normalized genes and one 60-gene pass per case, L1 ratio 0.7, four workers, seed 42, gap target 0.10, and 300-second limit. Final-figure evaluation independently mapped groups to their most frequent reference population, resolving ties by sorted reference name and permitting multiple groups per population. Macro-F1 averages equally across all four populations.

## Availability of Data and Materials

### Data Availability

The accessibility details of the 8cube dataset are described in the Data Availability section of (27). The MERFISH data are available through the Allen Brain Cell Atlas, under dataset `Zhuang-ABCA-2`. We used the isocortical cells from section `Zhuang-ABCA-2.033`. The Zhengmix4eq dataset is available through the Bioconductor package DuoClustering2018, using the function `sce_full_Zhengmix4eq()`.

### Code availability

CellSieve python package and documentation: https://github.com/pachterlab/CellSieve. Code to regenerate figures and analyses https://github.com/pachterlab/SP_2026. Data to reproduce the figures and analyses:.

## Competing Interests

The authors declare that they have no competing interests.

## Funding

This work was funded in part by grant NIH UM1HG012077.

## Author contributions

N.P.S. contributed to conceptualization, performed analyses, developed the CellSieve python package, made figures, and drafted the manuscript. L.P. contributed to conceptualization, supervision, and funding acquisition. All authors contributed to manuscript review.

## Acknowledgments

We thank the IGVF consortium for valuable discussion and feedback.

## Generative AI statement

We used OpenAI Codex to assist with code refactoring and development of reproducible analysis and figure-generation scripts. OpenAI ChatGPT was also used to assist with initial drafting and language refinement of portions of the manuscript. All AI-assisted text was subsequently substantially revised and rewritten by the authors, who take full responsibility for the final content. All resulting code, analyses, figures, and manuscript text were reviewed and validated by the authors.

## Notes

### Competing Interest Statement

The authors have declared no competing interest.

