## Supplementary Figures for "Gene set selection for classification with single-cell RNA-seq"

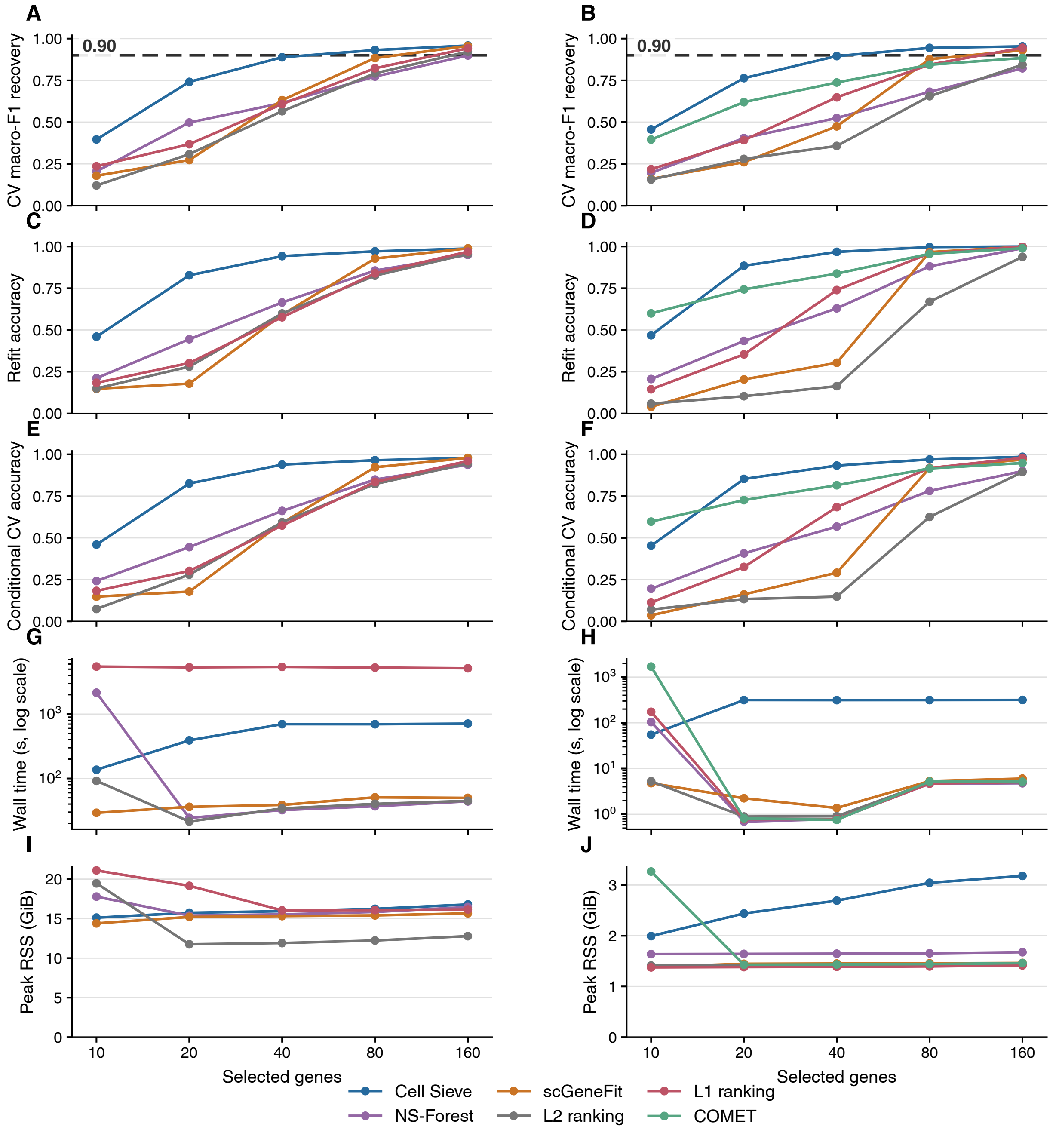


Figure S1: **Cortex classification performance and computational cost.** Full cortex/hippocampus (left) and the shared stratified subset (right). A–B, conditional CV macro-F1; C–D, refit accuracy; E–F, conditional CV accuracy; G–H, wall-clock time; I–J, peak process-tree memory.

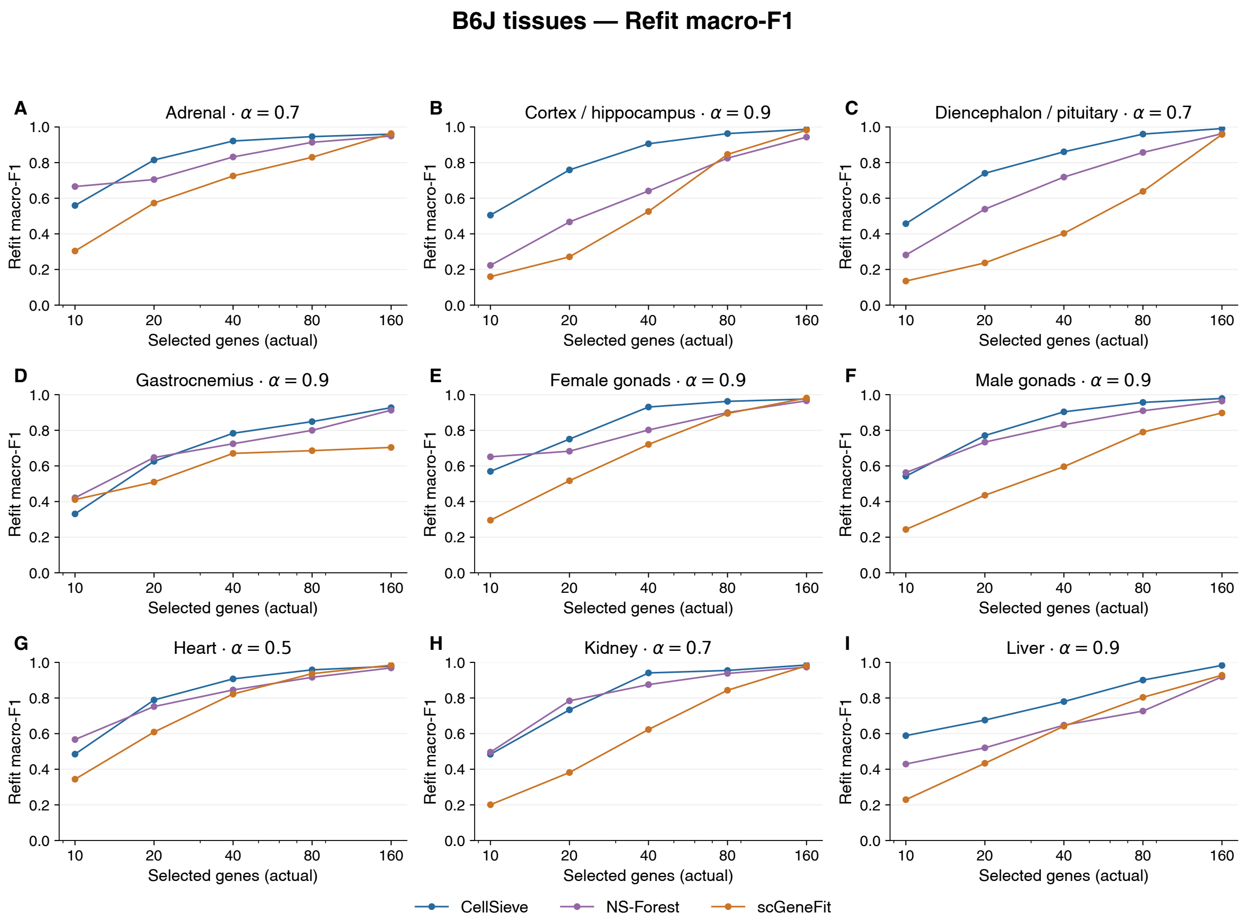


Figure S2: **Gene-panel performance across nine B6J tissues.** Refit macro-F1 for CellSieve, NS-Forest, and scGeneFit across panel sizes. Tissues and tissue-specific L1 ratios are indicated above each panel.

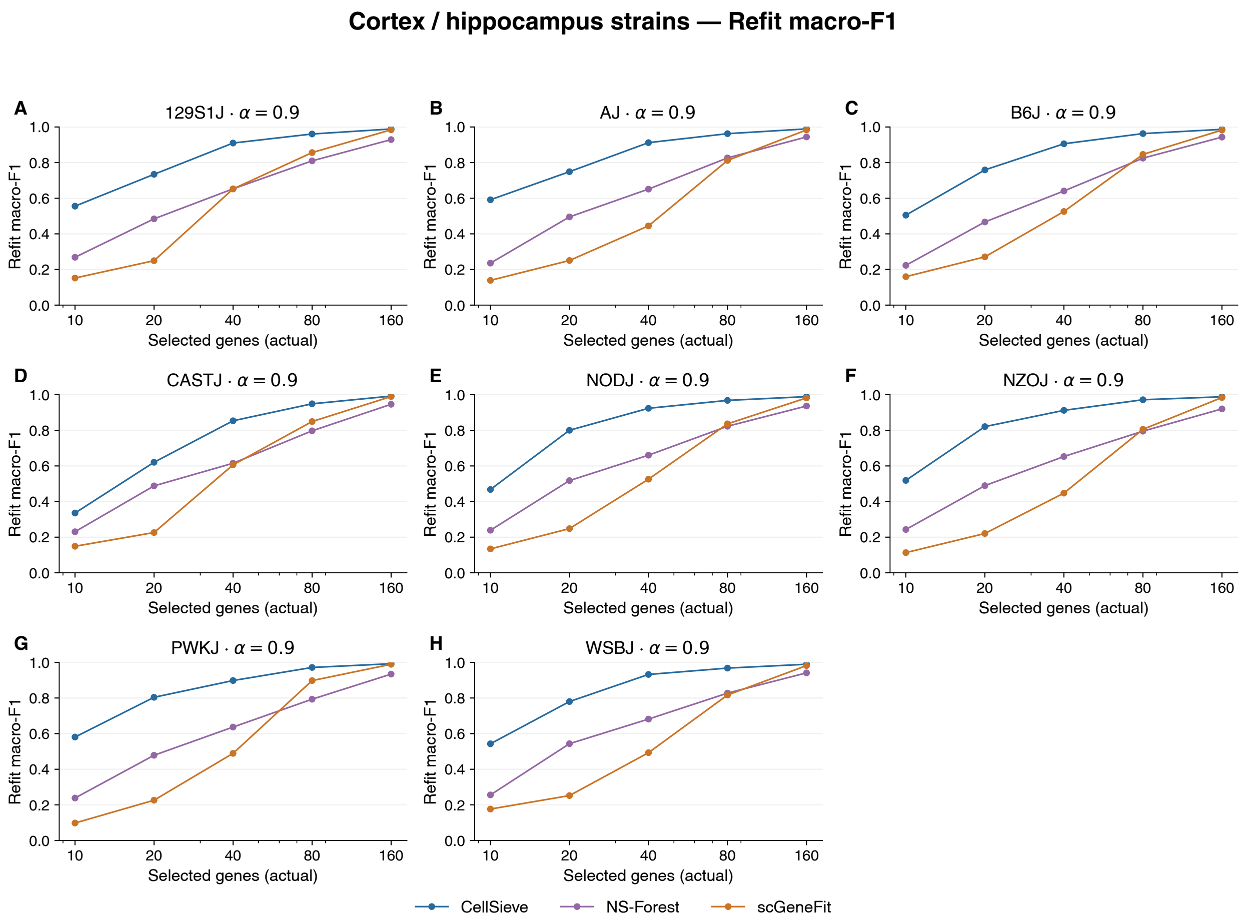


Figure S3: **Gene-panel performance across eight founder strains.** Refit macro-F1 for CellSieve, NS-Forest, and scGeneFit in cortex/hippocampus across panel sizes. CellSieve uses an L1 ratio of 0.9 for all strains.

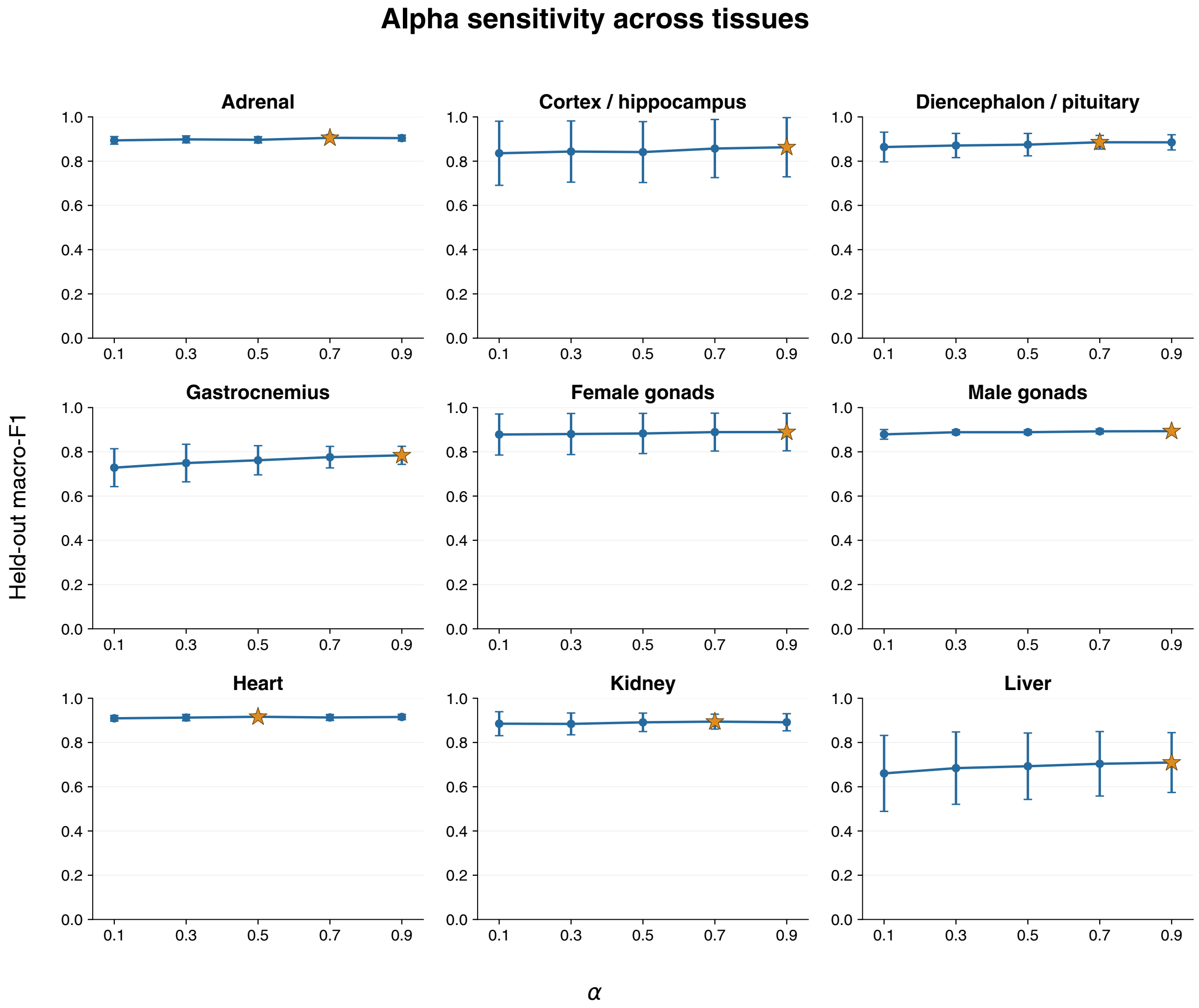


Figure S4: **Sensitivity to the elastic-net L1 ratio.** Held-out macro-F1 for 40-gene panels across nine B6J tissues. Points and error bars show mean and standard deviation over 20 paired subsamples. Stars mark the highest observed mean, not statistical significance.
